# Crotamine as a vehicle for non-viral gene delivery for Pompe disease

**DOI:** 10.1101/2021.03.23.436632

**Authors:** Frank Martiniuk, Adra Mack, Justin Martiniuk, Richard Karpel, Peter Meinke, Benedikt Schoser, Feng Wu, Kam-Meng Tchou-Wong

**Author notes:** 1 to whom reprints should be directed. shared equally in authorship and experiments (Adra Mack, Justin Martiniuk).

## Abstract

Genetic deficiency of lysosomal acid alpha glucosidase or acid maltase (GAA) results in Pompe disease (PD), encompassing at least five clinical subtypes of varying severity. The current approved enzyme replacement therapy (ERT) for PD is via IV infusion every 2 weeks of a recombinant human GAA (rhGAA) secreted by Chinese hamster ovary (CHO) cells (alglucosidase alfa/Myozyme, Sanofi/Genzyme). Although alglucosidase alfa has proven to be efficient in rescuing cardiac abnormalities and extending the life span of the infantile form, the response in skeletal muscle is variable. ERT usually begins when the patients are symptomatic and secondary problems are already present which are compounded by low alglucosidase alfa uptake, transient nature (every 2 weeks with a rapid return to defect levels), variable glycogen reduction, autophagic accumulation, immune response and high cost. A consensus at a recent US Acid Maltase Deficiency (AMD) conference suggested that a multi-pronged approach including gene therapy, diet, exercise, etc. must be evaluated for a successful treatment of PD. Compared to replication defective viruses, non-viral gene transfer offers fewer safety concerns and, if recent studies are validated, has a wider range of cells. In order for gene therapy (GT) to succeed, the gene of interest must be delivered into the affected cell and expressed to overcome the inherited deficiency. Cell penetrating peptides (CPPs) enter eukaryotic cells through an energy-independent mechanism and efficiently carry biologically active and therapeutic molecules into cells and localize in the cytoplasm or nucleus. CPPs are usually covalently linked to the cargo, including peptides and DNA. Crotamine (Cro) from the South American rattlesnake-*Crotalus durrissus terrificus* venom, can bind electrostatically to plasmid DNA to deliver into cells, including muscle. We have assembled a bacterial expression vector for Cro and purified the recombinant Cro (rCro). Transient transfection in AMD fibroblasts and *ex vivo* in whole blood from an adult Pompe patient with rCro complexed with the pcDNA3 x *hGAA* cDNA demonstrated increased GAA activity. In GAA knockout (KO) mice receiving a single injection of rCro complexed to pcDNA3 x *hGAA* cDNA intraperitoneally (IP), we found increased GAA activity in tissues after 48 hr. After 8 treatments-IP over 55 days, we found increased vertical hang-time activity, reduced glycogen deposition, increased GAA activity/*hGAA* plasmid in tissues and minimal immune-reaction to rCro. A subsequent study of 5 administrations every 2 to 3 weeks showed reverse of the clinical phenotypes by running wheel activity, Rotarod, grip-strength meter, open field mobility and T-maze. Tissue culture experiments in PD fibroblast, lymphoid and skeletal muscle cell lines showed increased GAA activity after rCro transient gene delivery.

## Introduction

Lysosomal acid maltase or GAA hydrolyzes linear α-1,4 glucosidic linkages in substrates ranging from large polymers (glycogen) to maltose and the artificial substrate 4-methylumbelliferyl-α-D-glucoside (1-18). Genetic deficiency of GAA results in Pompe disease (PD) encompassing at least 5 subtypes of varying severity (1,19) presenting as intracellular accumulation of glycogen in multiple tissues with skeletal muscle being the primary target, manifesting as myopathy and cardiomyopathy. The infantile PD is characterized by no GAA activity, muscle hypotonia, feeding difficulties and hypertrophic cardiomyopathy leading to death within the first year of life (20). The juvenile and adult onset forms are caused by partial enzyme deficiency, manifesting with progressive muscle weakness leading to wheelchair/ventilator dependence and premature death from respiratory insufficiency (21-24). The current approved enzyme replacement therapy (ERT) for PD is via IV infusion every 2 weeks of a recombinant human GAA (rhGAA) secreted by CHO cells (alglucosidase alfa/Myozyme, Sanofi/Genzyme). Although alglucosidase alfa has proven to be efficient in rescuing cardiac abnormalities and extending the life span of the infantile form, the response in skeletal muscle is variable. In juvenile and adult onset patients, only mild improvements in motor and respiratory functions have been achieved and is unsatisfactory in the reversal of skeletal muscle pathology. In infants, muscle pathology and degree of glycogen deposition correlates with the severity of symptoms and the earlier ERT is introduced, the better chance of response (24). In infants, glycogen is found mostly within large neurons of hindbrain, spinal cord and sensory ganglia with severe degeneration in axon terminals of primary sensory neurons (25-27). Treatments must address the neuropathology and hearing loss in infants (28-35). In juvenile and adult onset patients, mild improvements in motor and respiratory functions have been achieved, but are unsatisfactory in the reversal of skeletal muscle pathology (1,20-24, 36-40). Lysosomal glycogen accumulation leads to multiple secondary abnormalities (autophagy, lipofuscin, mitochondria, trafficking and signaling) that may be amenable to long-term therapy. ERT usually begins when the patients are symptomatic and secondary problems present (41). Gutschmidt et al. (42) found that ERT in late onset patients’ clinical outcomes, particularly lung function, muscle strength, and walking capability tend to deteriorate over time, indicating that additional efforts must be made to improve alglucosidase alfa’s effectiveness and supplement therapies developed. Some patients experienced infusion-associated reactions due to bronchial spasm with flushing and pulmonary artery blockage during infusion. Although alglucosidase alfa has been a wonderful first step in treating PD, it has revealed subtle aspects that must be dealt with for successful treatment (2,43-49). A consensus at a recent US Acid Maltase Deficiency (AMD) conference in San Antonio, Texas, suggested that a multi-pronged approach including gene therapy, diet, exercise, etc. must be evaluated for a successful treatment of PD.

Compared to replication defective viruses, non-viral GT offers fewer safety concerns and, if recent studies are validated, has a wider range of cells. Dunbar et al. (50) reviewed the current status of gene therapies with viral vectors, adoptive transfer of genetically engineered T-cells, hematopoietic stem cells and gene editing after 30 years of promise and setbacks. GT is rapidly becoming a critical component for inherited immune disorders, hemophilia, eye/neurodegenerative disorders and lymphoid cancers. The CRISPR/Cas9 gene editing approach would provide ways to correct or alter an individual’s genome with precision. Non-viral GT and their products should reduce cost and minimize immune responses and toxicity.

In order for GT to succeed the gene of interest must be delivered into the affected cell and expressed to overcome the inherited deficiency. Cell penetrating peptides (CPPs) have the ability to enter eukaryotic cells through an energy-independent mechanism and can carry biologically active and therapeutic compounds. CPPs are usually multiple sequences of short and positively charged peptides, rich in arginine and lysine residues, which penetrate cellular membranes and localize in the cytoplasm or the nucleus (51-53). CPPs are usually covalently linked to the cargo or synthesized as a fusion protein (54,55) including peptides/proteins, DNA, siRNA and liposomes (51,53,55,56). Studies suggest CPPs translocate cells via endocytosis (57), non-endocytotic pathways (58,59), pore formation (57,60) and membrane-associated heparan sulfate (61-72). Crotamine (vCro) is a myonecrotic polypeptide in the venom of the S. American rattlesnake, *Crotalus durissus terrifcus*. Radis-Baptista et al. (73) isolated the 340 bp *crotamine* cDNA from venom glands. It contained an open reading frame of 198 bp with 5’-50 and 3’-30 bp untranslated regions, a signal peptide sequence and a poly (A+) signal. vCro is a 42 amino acid cationic polypeptide with 11-basic residues and 6-Cys forming 3 disulfide bonds in -YKQCHKKGGHCFPKEKICLPPSSDFGKMDCRWRWKCCKKGS-G. vCro contains an antiparallel β-sheet (residues 9–13 and 34–38) and an α-helix (residues 1–7) stabilized by 3 disulfide bonds between Cys4-Cys36, Cys11-Cys30 and Cys18-Cys37 (74). Kerkis et al. (75) reported vCro can penetrate into different cell types, specifically associates with centrosomes/chromatin in the nucleus, binds to the chromosomes at the S/G2 phase and is stable over a wide pH range and temperatures. Nascimento et al. (76) showed that vCro is unique by binding electrostatically to plasmid DNA. Two to 4 µg of DNA complexed with 10 µg of vCro was effective in transfecting cultured cells. Mice treated-IP with 2 µg of DNA complexed with 10 µg of vCro for 24 hr or 30 days showed bone marrow, liver, bronchioles/lung, spleen, muscle, heart and brain were transduced (77-87). vCro does not demonstrate any immune-toxicity after long-term treatment (daily-IP of 1 μg for 21 days) in mice (78,88,89). Proteomics showed that small snake venom proteins of <5kD as vCro do not produce antibody response (90-92). Yamane et al. (93) showed specificity for proliferating cells, low cytotoxic effects and no hemolytic activity of vCro against non-tumor mammal cell lines. Chen et al. (94) demonstrated in a 3D model of vCro that residues 31-Arg-Trp-Arg-Trp-Lys-35 are potential DNA binding sites. Porta et al. (95) found that chemically synthesized full-length Cro compared favorably with its native counterpart including the ability to transfect and transport DNA into eukaryotic cells.

In this study, we have assembled a bacterial expression vector for Cro and purified the rCro. Transient transfection in PD transformed fibroblast cells and *ex vivo* in whole blood from an adult patient with rCro complexed with the pcDNA3 x *hGAA* cDNA demonstrated increased GAA activity. In GAA KO mice receiving a single injection of rCro x pcDNA3 x *hGAA* cDNA IP, we found increased GAA activity in tissues after 48 hr. After 8 treatments-IP over 55 days, we found increased vertical hang-time activity, reduced glycogen deposition, increased GAA activity/*hGAA* plasmid in tissues and minimal immune-reaction to rCro. A subsequent study of 5 administrations showed reverse of the clinical phenotypes by running wheel activity, Rotarod, grip-strength meter, open field mobility and T-maze. Experiments in PD fibroblast, lymphoid lines and skeletal muscle cell lines showed increased GAA activity after rCro transient gene delivery.

## Materials and Methods

### Construction of a Cro prokaryotic expression vector and expression/purification in E. coli

We constructed a prokaryotic expression mini-gene with the *Cro* cDNA in pUC19 under the control of the *LacZ* promoter/operator with a 5’-His tag and a Xa protease site (96,97). rCro was induced with IPTG for ∼2 hr and purified by nickel affinity chromatography via the His-tag, followed by Xa protease digestion (Novagen, Inc.) and analyzed by SDS-PAGE.

### Cell culture and transfection of PD human skeletal muscle, fibroblast and lymphoid cell lines

Human lymphoid (GM6314, GM13793, GM14450) or fibroblast (GM4912, GM1935, GM3329) or transformed TR4912 cell lines from infantile or adult PD were maintained in 15% fetal bovine serum, RPMI 1640 or DMEM supplemented with glutamine, penicillin and streptomycin at 37°C-5% CO_2_. A human PD skeletal muscle cell line (homozygous for the IVS1 c.-32– 13t>g)(98) and normal skeletal muscle cells were grown in PromoCell skeletal muscle growth media. Cells were plated at 0.3-0.4 × 10^6^ in 1.5 ml media in 6-well plates-24 hours before transfection with either rCro or lipofectamine 3000 (ThermoFisher) or Lipofectin (99-102) with the mammalian expression plasmid-pcDNA3 x human *GAA* cDNA (bp 1 to 3,150, 2,856 bp of coding cDNA (10,99-104). The rCro-*GAA* plasmid complexes were formed by mixing 1, 2 or 4 µg of plasmid DNA with 10 µg of rCro in 150 mM NaCl or PBS at RT for 10 min. Cells were incubated for 48 hr and assayed for GAA with 4-methylumbelliferyl-α-D-glucoside (4-MU-Glyc) at pH 4 and as an internal control, neutral α-glucosidase (NAG)(see below). Cells were harvested after 48 hr, washed with PBS, lysed by addition of 0.5 ml of 0.01 M sodium phosphate pH 7.5, frozen and thawed 2-3x, spun for 5 minutes in a microfuge to clarify and assayed for human GAA and NAG as described below. Cells were also exposed to a rhGAA (R and D Systems #8329-GH-025).

### *Ex vivo* studies

We transfected 3 x 3ml heparinized whole blood from an adult PD patient with 4 µg of the *hGAA* cDNA plasmid complexed with 10 µg of rCro or Lipofectin, incubated at 37°C for 24 hrs and WBCs isolated by hypaque-ficol density centrifugation. Cells were assayed for GAA/NAG as below.

### Enzyme Assay

The lysate (50 μl) was assayed for GAA in 100 μl of the artificial substrate 4-methylumbelliferyl-α-D-glucoside (4-MU-Glyc) (one mg per ml) in 0.5 M sodium acetate pH 4.0 for 6 to 18 hours at 37oC and fluorescence determined in a fluorometer (excitation-360_nm_ and emission-460_nm_) (Sequoia-Turner, Mountainview, CA). For assay of neutral α-glucosidase (NAG) as an internal control, 4-MU-Glyc in 0.5 M sodium phosphate at pH 7.5 was used (10,105). All assays were done in triplicate. Bradford protein assay used 150 μl reagent (BioRad #500-0006) diluted 1->5 with water in a 96 well flat-bottomed microtiter plate. Standards were serial dilutions of BSA from 1000 μg/ml to 2 μg/ml for 1 to 25 μl sample volume. Samples were read at A_595_nm before 60 min.

### In vivo studies GAA KO mice

We utilized the GAA KO mouse developed by Raben et al. (106) with the exon 6_neo_ disruption (Jackson Labs; B6,129-Gaa^tm1Rabn^/J, exon 6_neo_), wild-type BALB/c or 129/J or GAA KO mice mock-treated with PBS. We IP infused GAA KO mice (∼4-6 months old) with a single or multiple doses of 4 or 20 μg of *hGAA* plasmid complexed to 10 μg rCro. At 7 days, mice were sacrificed and tissues were assayed for GAA and NAG and compared to wild-type mice and mock treated GAA KO mice. Also, a weekly treatment for 55 days and 8 treatments with 10 μg rCro complexed with 0 or 4 μg or 20 μg *hGAA* plasmid and measured vertical hang-time (forelimb strength)(106), GAA activity/plasmid content in tissues, immune-reaction to Cro and glycogen deposition.

### Phenotypic evaluation of GAA KO mice after rCro-hGAA plasmid treatment

GAA KO mice (106) were treated IP with either 4 µg or 10 µg of *hGAA* plasmid complexed to 10 µg of rCro in 100 μl PBS every 2 to 3 weeks for 5 administrations. Tissues, urine, blood slides, weight and serum were collected. Mice were tested every ∼3 weeks between 8:00-11:00AM to avoid circadian variation for: motor activity by running wheel (RW-Mini-Mitter Co., Inc., OR); fore-limb muscle strength by grip-strength meter (GSM)(Columbus Inst. OH), motor coordination/balance with a Rotarod (AJATPA Expert Industries, India), open-field mobility by 5 min video/distance traveled and spatial preference/spontaneous learning with a T-maze (Stoelting, IL) (100-102).

### ELISA for antibodies to rCro

In Falcon plates #3912, rCro was added at 1 μl/well in 100 μl PBS at 4°C overnight, washed 3x with PBS and blocked with 200 μl PBS-1g% BSA-0.05% Tween-20 for 18 hr at RT. Then, add 1 μl serum in 100 μl PBS-overnight at 4°C, wash 3x with PBS, 1x-PBS-0.05% Tween-20 and 2X PBS. Add 100 μl goat anti-mouse IgG-HRP (BioRad #172-1101) at 1:500 in PBS/BSA/Tween-20 at 4°C overnight. Wash 3x PBS, 1x PBS-0.05% Tween-20, 2x PBS and stain with 100 μl of 3,3’,5,5’–tetramethylbenzidine (Sigma #T5525) in 9 ml of 0.05 M phosphate-citrate buffer, pH 5.0 (Sigma #P4922) with 20 μl of 30% H_2_O_2_ for 30-60 minutes at room temperature. The reaction is stopped with 50 μl 1 N HCl and read at A_450_nm

### Assay for glycogen content

In a 96 well microtiter plate, add 25 μl sample, 25 μl 0.05 M sodium phosphate pH 6.5, +/- 0.5 μl amyloglucosidase (Sigma #A-1602), placed overnight at 37°C (107). Glycogen standards (rabbit liver, 2 mg/ml) and D-Glucose standards were at 400 μM, 200 μM and 100 μM. Add 20 μl of Eton Bioscience glucose assay solution (#1200031002) and incubate for 30 minutes at 37°C. The reaction was stopped with 25 μl of 0.5 M acetic acid and read at A_490_nm.

### PCR for hGAA plasmid in tissues

PCR utilized Bulldog Fastmix Frenchie according to instructions with 1 μg genomic DNA from tissues after 8 treatments or 55 days or 1 ng of pcDNA3x*hGAA* cDNA. DNA was extracted using standard phenol-chloroform methods. The human *GAA* cDNA primers are: GAAFM106 5’-CCATTTGTGATCTCCCGCTCGACCTTTG-3’ (bp1733-1810) and GAAFM94 5’-GGTGGTCCACAGTCCAGGTGCTAGAG-3’ (actual primer sequence; bp2201-2227). The cycling conditions are: hot-start 94oC for 2 min, disassociation-94oC for 30 sec, annealing at 58oC for 30 sec and extension at 72oC for 1 min for 40 cycles with a final extension for 10 min. The PCR amplicons (20 μl) are electrophoresed in 1.5% agarose and visualized under UV light with ethidium bromide staining.

### Statistical analysis

Student t-test for probability associated with a population and standard deviation based upon the population were determined using Microsoft Excel software and considered significant at p value ≤ 0.05.

## Results

### Construction of an rCro expression vector and expression/purification in *E. coli*

We constructed a prokaryotic expression mini-gene with the *Cro* cDNA in pUC19 under the control of the *LacZ* promoter/operator with a 5’-His tag and a Xa protease site (96,97). The rCro was induction with IPTG for ∼2hr and purified by nickel affinity chromatography via the His-tag, followed by Xa protease digestion and analyzed by SDS-PAGE (76,78). **Figure 1** shows in lane 1-lysate pre-Ni column; lane 2-lysate post Ni column; lane-3 first wash fraction of Ni column and lane 4-eluted fraction with the arrow showing the correct size of rCro at ∼5 kD. The rCro was also characterized for binding time to plasmid DNA and plasmid binding capacity. The rCro showed maximum plasmid binding of 4 µg to 10 µg rCro and a minimum time of one minute similar to the published data.

**Figure 1.**
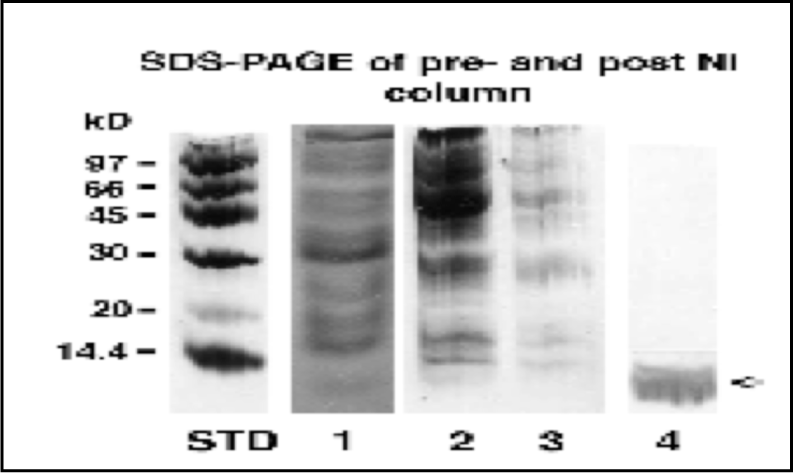
Composite diagram of a rCro prokaryotic expression vector and expression/purification in *E. coli*. SDS-PAGE shows standards in the left lane, lane 1-lysate pre-Ni column; 2-lysate post Ni column; 3-first wash fraction of Ni column and 4-eluted fraction with the arrow showing the correct size of rCro at ∼5kD.

### rCro transient transfection of an PD transformed fibroblast cell line

TR4912, an immortalized infantile PD fibroblast cells (no GAA activity or mRNA)(108) were transfected with various amounts (1, 2 or 4 µg) of *hGAA* plasmid complexed to 10 µg of rCro. After 48 hr, cells were assayed for GAA and NAG (**Graph 1**). As a positive control for transfection, cells were transfected with Lipofectin and *hGAA* plasmid. Transfected TR4912 cells increased from 0.0001 to 5, 16 and 48 GAA/NAG ratio for 1, 2 or 4 μg plasmid respectively with 10 μg rCro. Cells transfected with Lipofectin (99-102) had a ratio of 12, while all mock treated cells were undetectable. GAA/mg protein did not change results (not shown). Thus, rCro mediated gene delivery was greater than lipid mediated.

### *Ex vivo* studies

We transfected 3 x 3 ml heparinized whole blood from an adult PD patient with 4 µg of the *hGAA* cDNA plasmid complexed with 10 µg of rCro or Lipofectin that was incubated at 37°C for 24 hr and WBCs isolated by hypaque-ficol density centrifugation. Cells were assayed for GAA/NAG ratio. Mocked treated WBCs had a relative GAA/NAG ratio of 5; blood treated with the *hGAA* plasmid x rCro complexes had a GAA/NAG ratio of 45, while lipid had a GAA/NAG ratio of 25.

### Short-term *in vivo* rCro gene delivery in GAA KO mice

4 μg of *hGAA* plasmid was complexed to 10 μg rCro that was administered IP to three GAA KO mice in 100 μl-PBS. After 48 hr, tissues were assayed. Tissues-heart, skeletal muscle, liver, kidney and diaphragm showed significant increases (**Table 1**)(mean±SD). GAA/mg protein did not change results (not shown).

**Table 1.**
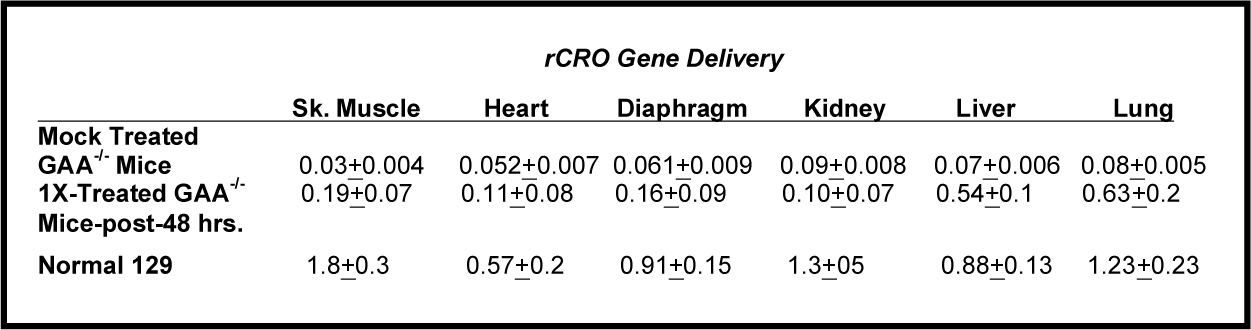

### Long-term *in vivo* rCro gene delivery and vertical hang-time measurement

We are utilizing non-viral *in vivo* gene delivery with rCro that is transient in nature as an episome (maximum expression for 3-14 days)(109-126), thus we decided upon a weekly treatment for 55 days or 8 treatments schedule to maintain and monitor sustained GAA levels. We treated four groups (age matched) of GAA KO mice-IP (n=2-3) weekly with 10 μg rCro complexed with 0 or 4 μg or 20 μg *hGAA* plasmid and measured vertical hang-time (forelimb strength)(106). The 8 times treated animals were well tolerated as mice gained weight over 55 days (data not shown). Both treated groups showed a steady and significant improvement in vertical hang-time (**Figure 2**). Interestingly, mice in the 4 μg plasmid group gradually increased to the 20 μg plasmid group, suggesting that a lower dose may be effective long-term. Both treated groups showed ∼67% of wild-type mice hang-time. All treatments were significant at p=≤0.05.

**Figure 2.**
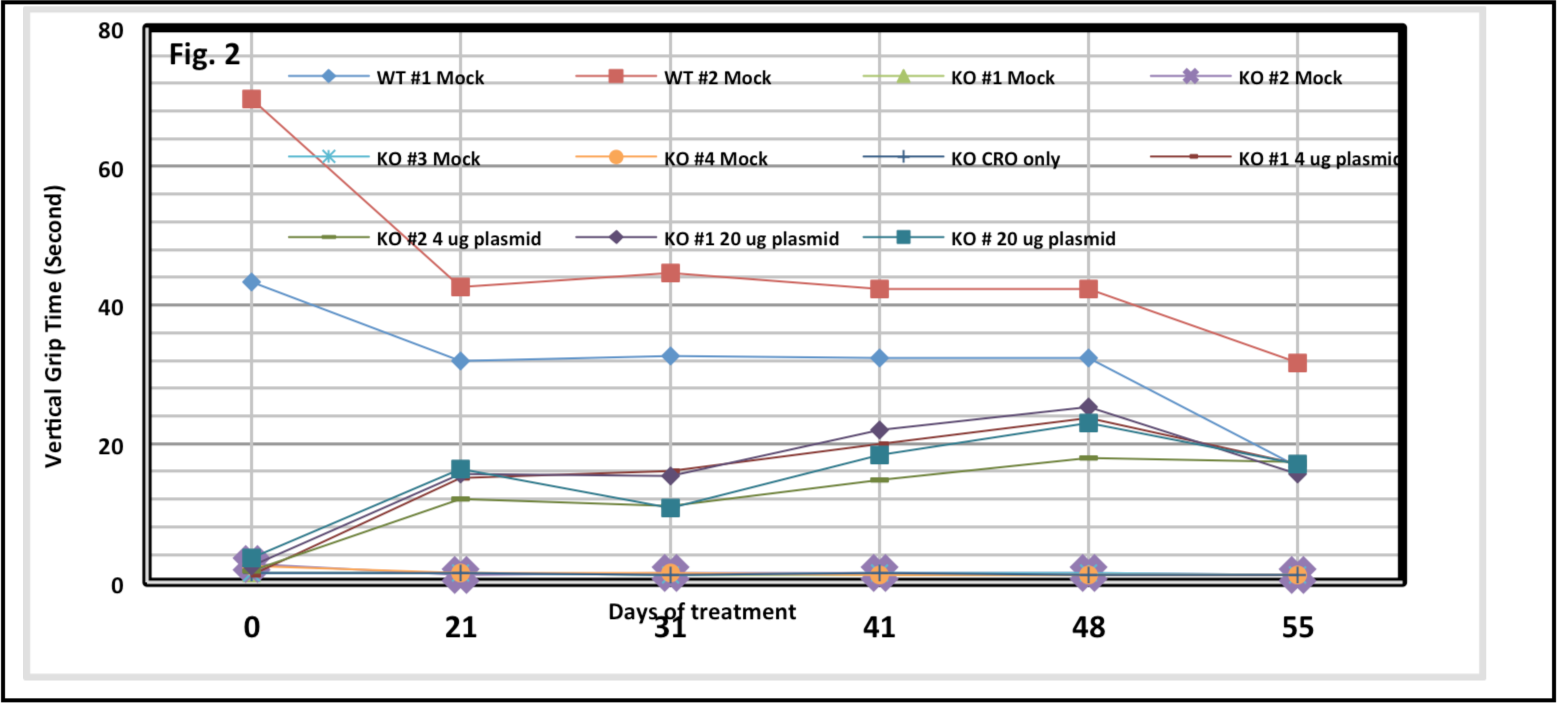
Long-term *in vivo* treatment and vertical hang-time measurement. We treated four groups (age matched) of GAA KO mice-IP (exon 6_neo_)(n=2-3) weekly with 10 μg rCro complexed with 0 or 4 μg or 20 μg *hGAA* plasmid and measured vertical hang-time (forelimb strength). Both treated groups showed a steady and significant improvement in vertical hang-time. Interestingly, mice in the 4 μg plasmid group gradually increased to the 20 μg plasmid group. Both Tx groups showed ∼67% of WT mice hang-time. Mean±SD not shown to not complicate the graph.

### Biodistribution (BD) in tissues and glycogen reduction after 8 treatments over 55 days

We found increased GAA activity in heart, hind-limb skeletal muscle, kidney, liver and diaphragm from treated GAA KO mice (**Table 2**) expressed as GAA/NAG. The % of wild type ranged from 11-32% (in red). Similar increases were found when expressed as GAA/mg protein (not shown). After 8 treatments over 55 days, we found tissue glycogen levels reduced from 11 to 96% (**Table 3**)(106). GAA/mg protein did not change results (not shown).

**TABLE 2.**

| TABLE 2. | <u>Mouse tissue GAA/NAG (mean ± SD) after 8 Tx over 55 days.</u> |  |  |  |  |
| --- | --- | --- | --- | --- | --- |
|  | Hind-limb Skeletal Muscle | Heart | Diaphragm | Liver | Kidney |
| Treated GAA KO-4ug plasmid + CRO | 0.22 ± 0.02 | 0.14 ± 0.02 | 0.18 ± 0.03 | 0.14 ± 0.04 | 0.23 ± 0.04 |
| % of WT | 28% | 18% | 28% | 11% | 21% |
| Treated GAA KO-20ug plasmid + CRO | 0.22 ± 0.01 | 0.11 ± 0.01 | 0.21 ± 0.04 | 0.21 ± 0.05 | 0.23 ± 0.01 |
| % of WT | 28% | 15% | 32% | 16% | 21% |
| Treated GAA KO-CRO | 0.054 ± 0.01 | 0.07 ± 0.01 | 0.077 ± 0.1 | 0.12 ± 0.08 | 0.29 ± 0.02 |
| Mock GAA KO-PBS | 0.07 ± 0.044 | 0.065 ± 0.044 | 0.08 ± 0.14 | 0.05 ± 0.009 | 0.15 ± 0.02 |
| Wild-type | 0.8 ± 0.11 | 0.76 ± 0.23 | 0.65 ± 0.38 | 1.33 ± 0.36 | 1.12 ± 0.08 |

**TABLE 3.**

| TABLE 3. | <u>% glycogen reduction (mean) compared to mock treated in GAA KO mouse tissues.</u> |  |  |  |  |
| --- | --- | --- | --- | --- | --- |
|  | Hind-limb Skeletal Muscle | Heart | Diaphragm | Liver | Kidney |
| Treated GAA KO -4ug plasmid + CRO | 96 | 11 | 47 | 29 | 48 |
| Treated GAA KO -20ug plasmid + CRO | 28 | 17 | 49 | 32 | 47 |

### Immune-antibody response

Serum antibody titer to rCro at 55 days/8 treatments for rCro alone was 35ng/ml, with rCro-4 μg plasmid = 70 ng/ml and for 20 μg plasmid = 60 ng/ml, thus demonstrating low immune response (78,87-92).

### PCR for *hGAA* plasmid in tissues

Agarose gel electrophoresis shows PCR amplicons from 1 μg genomic DNA from tissues (liver, heart, skeletal muscle, kidney and diaphragm) after 8 treatments over 55 days or positive control-1 ng of pcDNA3 x *hGAA* cDNA (**Figure 3**). We found the *hGAA* amplicon present in all treated tissues to varying degree, however the liver required nested PCR to detect. DNA from WT tissues (not shown) and mock treated GAA KO mice (lane 2) were negative for the *hGAA* amplicon.

**Figure 3.**
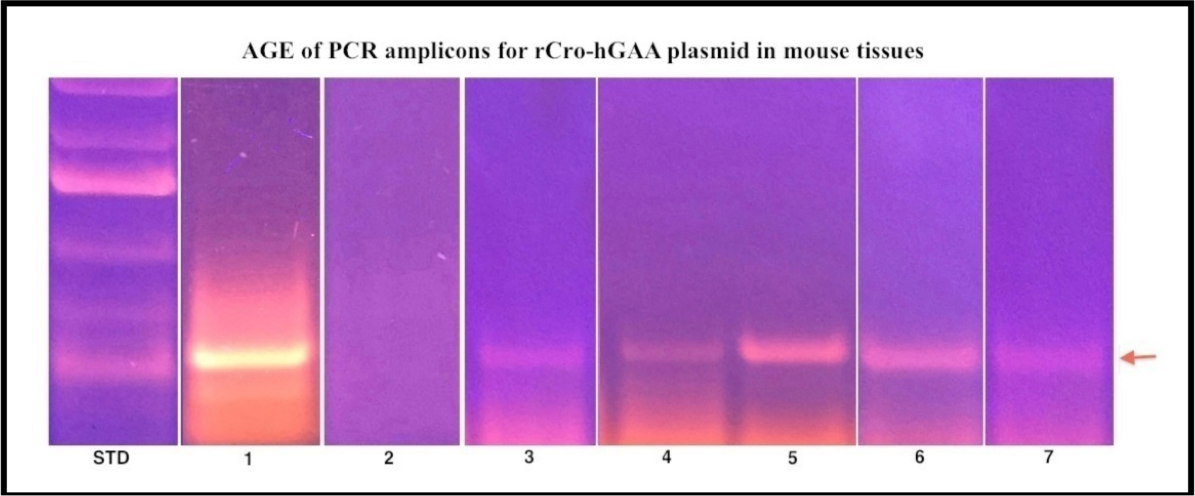
PCR for *hGAA* plasmid in tissues. PCR was performed on 1 μg genomic DNA from tissues after 8 treatments (Tx) or 1 ng of pcDNA3 x *hGAA* cDNA. Agarose gel electrophoresis of the amplicons-on left lane = STDs-ThermoFisher GeneRuler 1 kb ladder, lane 1=plasmid, lane 2=mock treated liver, lane 3=treated liver, lane 4=treated heart, lane 5=treated skeletal muscle, lane 6=treated kidney and lane 7=treated diaphragm.

### Assessment of spontaneous alternation in WT and GAA KO mice

Spontaneous alternation is used to assess the cognitive ability of rodents to choose one of the 2 goal arms of the T-maze. The advantage of a free choice procedure is that hippocampal or lesioned animals often develop a side preference and scores below 50%. Controls generally achieve at least 60-80% correct alternation. We assessed spontaneous alternative learning for cognitive ability in the T-maze in both male and female GAA KO mice and WT-129/C57 mice from 2-9 months of age. We found that deficiency in spontaneous learning appeared by 2-3 months in male and 3-4 months in female GAA KO mice (**Graph 2**). All conditions were significant (p=≤0.05).

### Phenotypic evaluation of GAA KO mice after rCro-*hGAA* plasmid treatment

GAA KO mice (106) were treated IP every 2 to 3 weeks with either 4 µg or 10 µg of *hGAA* plasmid complexed to 10 µg of rCro for 5 administrations (**Table 4**). Mice were tested every ∼3 weeks for: motor activity by running wheel (RW); fore-limb muscle strength by grip-strength meter (GSM), motor coordination/balance with a Rotarod, open-field mobility by 5 min video/distance traveled and spontaneous learning with a T-maze. All measurements improved significantly after 2 treatments for both doses of *hGAA* plasmid compared to pre-treatment measurements. All measurements were significant at p=<0.05 compared to pre-treated mice. All treatments were significant at p=≤0.05.

**Table 4.**

| Table 4 | RW | GSM | Rotarod | T-maze | Open-field mobility |
| --- | --- | --- | --- | --- | --- |
| Mean±SD | rev/12 hrs | lbs at release | Time (sec) | % correct alternation | cm/min ±SD |
| **GAA KO M/F** |  |  |  |  |  |
| Pre-Tx | 404±128 | 0.130±60 | 112±120 | 23±12 | 182±80 |
| **4ug plasmid/10ug Cro** |  |  |  |  |  |
| 2x injections | 2047±677* | 0.253±99* | 296±131* | 50±19* | 318±174* |
| 4x injections | 3156±647* | ND | ND | ND | ND |
| 5 x injections | 2177±647* | 0.232±87* | 287±145* | 51±7* | 229±73* |
| **10ug plasmid/10ug Cro** |  |  |  |  |  |
| 2x injections | 1730±366* | 0.293±117* | 131±59* | 58±13* | 388±86* |
| 4x injections | 3536±1687* | ND | ND | ND | ND |
| 5 x injections | 3508±2007* | 0.249±97* | 288±129* | 53±18* | 334±56* |
| **WT-129/C57 M/F n=8** | 4912±596 | 0.627±265 | 245±98 | 61±10 | 334±96 |

\*p<0.05 when compared to pre-treatment data by T-test (1-tail, 2-equal variance)

### rCro transient transfection in a human PD skeletal muscle, lymphoid and fibroblast cell lines

A human PD skeletal muscle cell line, three PD lymphoid and three PD fibroblast cell lines were transfected with 4 µg of *hGAA* plasmid complexed to 10 µg of rCro. After 48 hr, cells were assayed for GAA and NAG. Cells were also transfected with Lipofectamine 3000-*hGAA* plasmid or exposed to rhGAA (skeletal muscle only) for 48 hr and assayed. Mock treated PD and normal cell lines were controls. We found that lipo-*hGAA* plasmid increased GAA to 23% of normal while rCro-*hGAA* plasmid increased GAA to 28% of normal and rhGAA increased GAA to 20% of normal in PD skeletal muscle cells (mean+/SD)(**Graph 3**). In transfected PD fibroblast and lymphoid cell lines transfected with rCro-*hGAA* plasmid, increased GAA/NAG ratios was observed to various degrees as compared to normal cells (**Graphs 4** and **5**). All treatments were significant at p=≤0.05. GAA/mg protein did not change results (not shown).

## Discussion

Genetic deficiency of lysosomal GAA results in Pompe’s disease (PD), encompassing at least five clinical subtypes of varying severity (19). The current approved ERT for PD is via IV infusion every 2 weeks of a rhGAA secreted by CHO cells (alglucosidase alfa). Although alglucosidase alfa has proven to be efficient in rescuing cardiac abnormalities and extending the life span of the infantile form, the response in skeletal muscle is variable. In juvenile and adult patients, only mild improvements in motor and respiratory functions have been achieved and is unsatisfactory in the reversal of skeletal muscle pathology. ERT usually begins when the patients are symptomatic and secondary problems are already present which are compounded by low (about 1 %) alglucosidase alfa uptake in muscle (41), transient nature of administered (every 2 weeks with a rapid return to defect levels), variable glycogen reduction, autophagic accumulation, immune response and high cost. Although alglucosidase alfa has been a wonderful first step in treating PD, it has revealed subtle aspects that must be dealt with for successful treatment (2,43-49).

Compared to replication defective viruses, non-viral gene transfer offers fewer safety concerns and, if recent studies are validated, has a wider range of cells. Dunbar et al. (50) reviewed the current status of GTs with viral vectors, adoptive transfer of genetically engineered T-cells, hematopoietic stem cells and gene editing after 30 years of promise and setbacks. The CRISPR gene-editing approach would provide ways to correct or alter an individual’s genome with precision. Recent findings showed that there was low level of integration with AAV GT in dogs sometimes near genes affecting cell growth and must be monitored for development of cancers (127). There have been many studies using viral mediated GT in PD cells and animal models using adenovirus, adeno-associated virus, lentivirus and retroviruses (128-151). Barry Byrne and his colleagues at the Univ. of Florida and investigators at Duke University are developing GT using AAV viral systems (152-177). Over the next 10 years, viral delivered GTs are expected to come into their own as a treatment option for a variety of diseases with two therapies have regulatory approval (Novartis’ Zolgensma for spinal muscular atrophy and Spark’s Luxturna for a rare form of genetic blindness). GTs offer great rewards to treat many diseases, but there are also significant risks. Over the past year, several clinical studies have been placed on hold or stopped entirely due to safety concerns. Audentes Therapeutics had a temporary hold placed on the GT for X-linked myotubular myopathy following several patient deaths that has recently lifted by the FDA. Uniqure was placed on hold for its hemophilia B trial after a patient in the study developed liver cancer. Other studies found some patients developed acute myeloid leukemia (AML). A sickle cell disease clinical trial using CRISPR gene-editing to turn on a fetal form of hemoglobin reported promising results last year. However, CRISPR gene-editing can have off-target effects and rearrange chromosomes and if it can trigger cancer may not be known for several years. In December 2020, uniQure reported that a patient in a GT trial for hemophilia had developed a liver tumor and planned to evaluate whether its vector, an adeno-associated virus (AAV), had a role. Even though AAVs are supposed to be safer than lentiviruses because they are not designed to insert their gene or cargo into the genome, animal studies have found they can (178). While the precise mechanisms that led to these toxicities remain unknown, some hypotheses have focused on the role of antibodies to AAV that either preexist or rapidly accumulate following vector infusion. The formation of vector–antibody complexes could activate complement through the classic pathway or activate innate immunity through Fc-dependent uptake in antigen-presenting cells. The complexity of host–vector interactions in GT and the species-specific differences in those interactions have long limited the ability to use preclinical data to predict clinical outcomes. This is especially true with regard to immune-mediated vector toxicities. However, these limitations should not be seen as a reason to abandon animal modeling. Instead, the early clinical development of GT vectors may require an iterative process, especially if unexpected toxicities are observed (179-181).

Promising non-viral delivery systems are currently undergoing clinical trials (109-117,124-126). GT is the modulation of gene expression in specific cells to treat pathological conditions by introducing exogenous nucleic acids by carriers or vectors as DNA, mRNA, small interfering RNA (siRNA) or microRNAs. Conventional non-viral delivery methods are hydrodynamic pressure techniques, electroporation, ballistic bombardment and microinjection. Cationic polymer methods include lipofection, cationic peptides, PEI and receptor-mediated. We have previously investigated various delivery systems for non-viral GT with several mammalian expression plasmids containing the *hGAA* cDNA *in vitro* and *in vivo* were Helios gene gun (99,100), liposome (cationic lipid GL-67)(unpublished)(99-101) and PEI (unpublished) that included CMV, muscle specific and modified CMV regulatory promoters (101,102). In Australia with Peter Healy-DVM, two PD shorthorn cows were infused 4x over 4 months with a lipid-plasmid complex (150mg/100kg). Treated animals did not progress with muscle weakness and serum CPK dropped from 1000U pre-Tx to 500U by 4 months and were “healthy”. Skeletal muscle, diaphragm and intercostal muscle showed GAA increased from ∼1% to 10-20% of normal (unpublished).

In order for GT to succeed, the gene of interest must be delivered into the affected cell and expressed to overcome the inherited deficiency. CPPs enter eukaryotic cells through an energy-independent mechanism and efficiently carry biologically active and therapeutic molecules into cells and localize in the cytoplasm or nucleus. CPPs are usually covalently linked to the cargo, including peptides and DNA. vCro from the South American rattlesnake *Crotalus durrissus terrificus* venom can bind electrostatically to plasmid DNA to deliver into cells, including muscle. We decided to use a weekly administration regimen to obtain maximum and sustained GAA levels based upon the transient nature of non-viral GT and short-term episomal longevity. Similar to *in vitro* transfection, maximum expression and maintenance may be only 3-14 days before the plasmid is lost however long term detection has been observed (109-124). We have assembled a bacterial expression vector for Cro and purified the rCro. Transient transfection in PD fibroblasts and *ex vivo* in whole blood from an adult PD patient with rCro complexed with the pcDNA3 x *hGAA* cDNA, demonstrated increased GAA activity. In GAA KO mice receiving a single injection of rCro x *hGAA* cDNA IP, we found increased GAA activity in tissues after 48 hr. After 8 treatments-IP over 55 days, increased vertical hang-time activity, reduced glycogen deposition, increased GAA activity/*hGAA* plasmid in tissues and minimal immune-reaction to rCro. A subsequent study of 5 administrations showed reverse of the clinical phenotypes by running wheel activity, Rotarod, grip-strength meter, open field mobility and T-maze. Experiments in PD fibroblast, lymphoid lines and skeletal muscle cell lines showed increased GAA activity after rCro transient gene delivery.

From DNA vaccine (182-185), the production of the recombinant protein may be released into the circulation and taken up by distant tissues, functioning as an “enzyme-factory or depot.” With future experiments, if we achieve 10% of normal, we will consider the treatment a success since adult PD patients have 2-10% and survive to the 5/6^th^ decade. If over-expression is observed, we will add a Tet-off doxycycline (TRE) to regulate expression dose-dependently. If plasmid related cytotoxicity is found, we will remove CpG dimer sequences by minicircle synthesis (182,185). We also hypothesize there will be positive selective pressure to maintain the plasmid in tissues to be “healthier” by being able to hydrolyze glycogen to glucose (109-126,149-151). In some studies, plasmid integration into chromosomes was observed, however integration has the potential to cause tumorigenic events (109-126,149-151). Additionally, to further minimized immune-reaction to rCro, we will evaluate a smaller rCro (94).

## Acknowledgements

None

---

**Graph 1.**
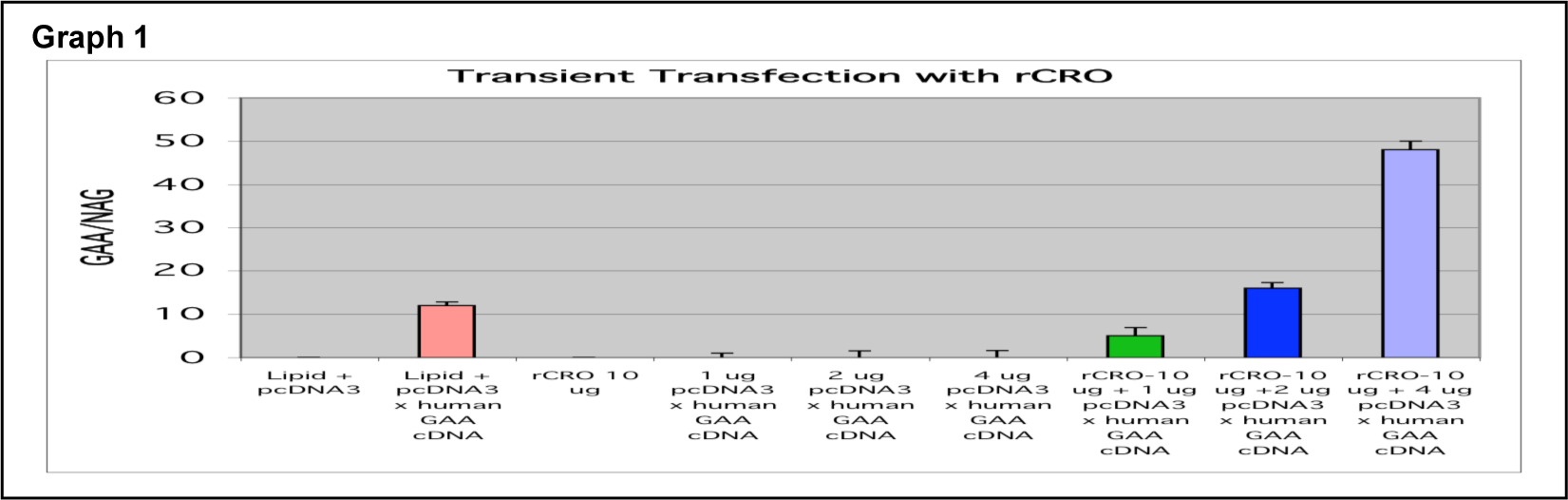
rCro transient transfection of a PD transformed fibroblast cell line. TR4912, an immortalized infantile PD fibroblast cells (no GAA activity or mRNA) were transiently transfected with rCro-*hGAA* plasmid complexes formed by mixing 1, 2 or 4 µg of plasmid DNA with 10 µg of rCro. Cells were incubated for 48 hr and assayed for GAA with 4-MU-Glyc at pH4 and as an internal control, neutral α-glucosidase (NAG). As a positive control for transfection, cells were transfected with Lipofectin and plasmid. Transfected TR4912 cells increased from 0.0001 to 5, 16 and 48 GAA/NAG for 1, 2 or 4 μg plasmid with 10 μg rCro. Cells transfected with Lipofectin had a ratio of 12, while all mock treated cells were undetectable.

**Graph 2.**
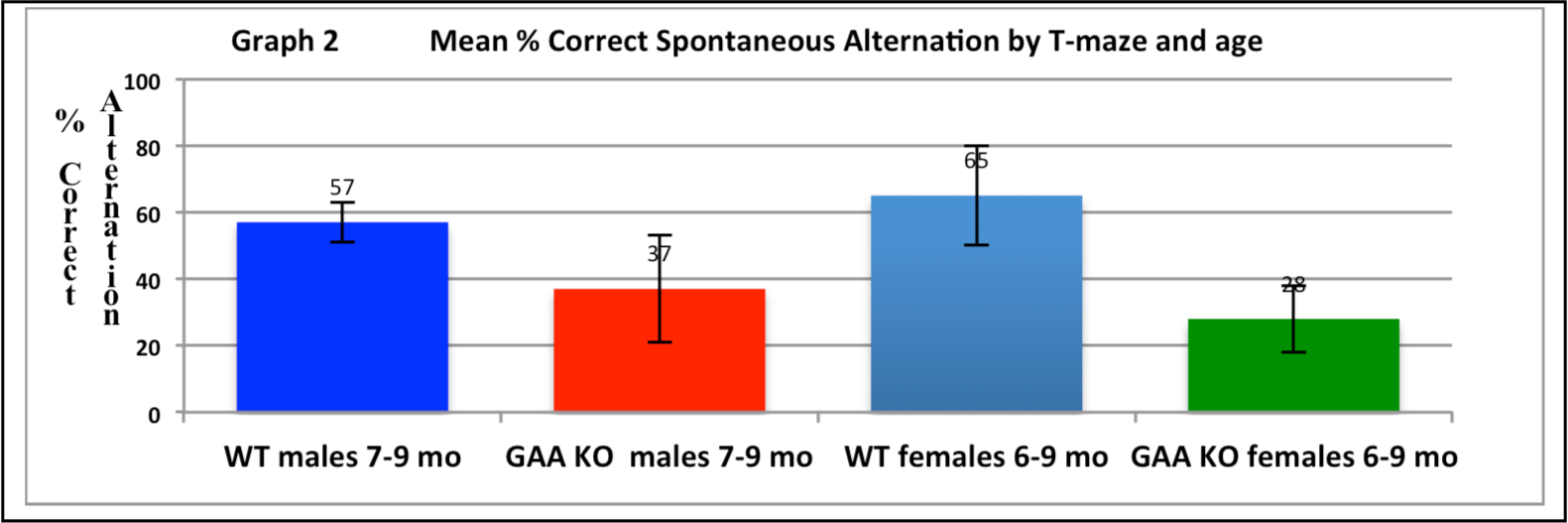
Mean % correct spontaneous alternation by T-maze and age. Spontaneous alternation is used to assess the cognitive ability of rodents to choose one of the 2 goal arms of the T-maze. The advantage of a free choice procedure is that hippocampal or lesioned animals often develop a side preference and scores below 50%. Controls generally achieve at least 60-80% correct alternation. We assessed spontaneous alternative learning for cognitive ability in the T-maze in both male and female GAA KO mice and WT-129/C57 mice from 2-9 months of age. All conditions were significant (p=≤0.05).

**Graph 3.**
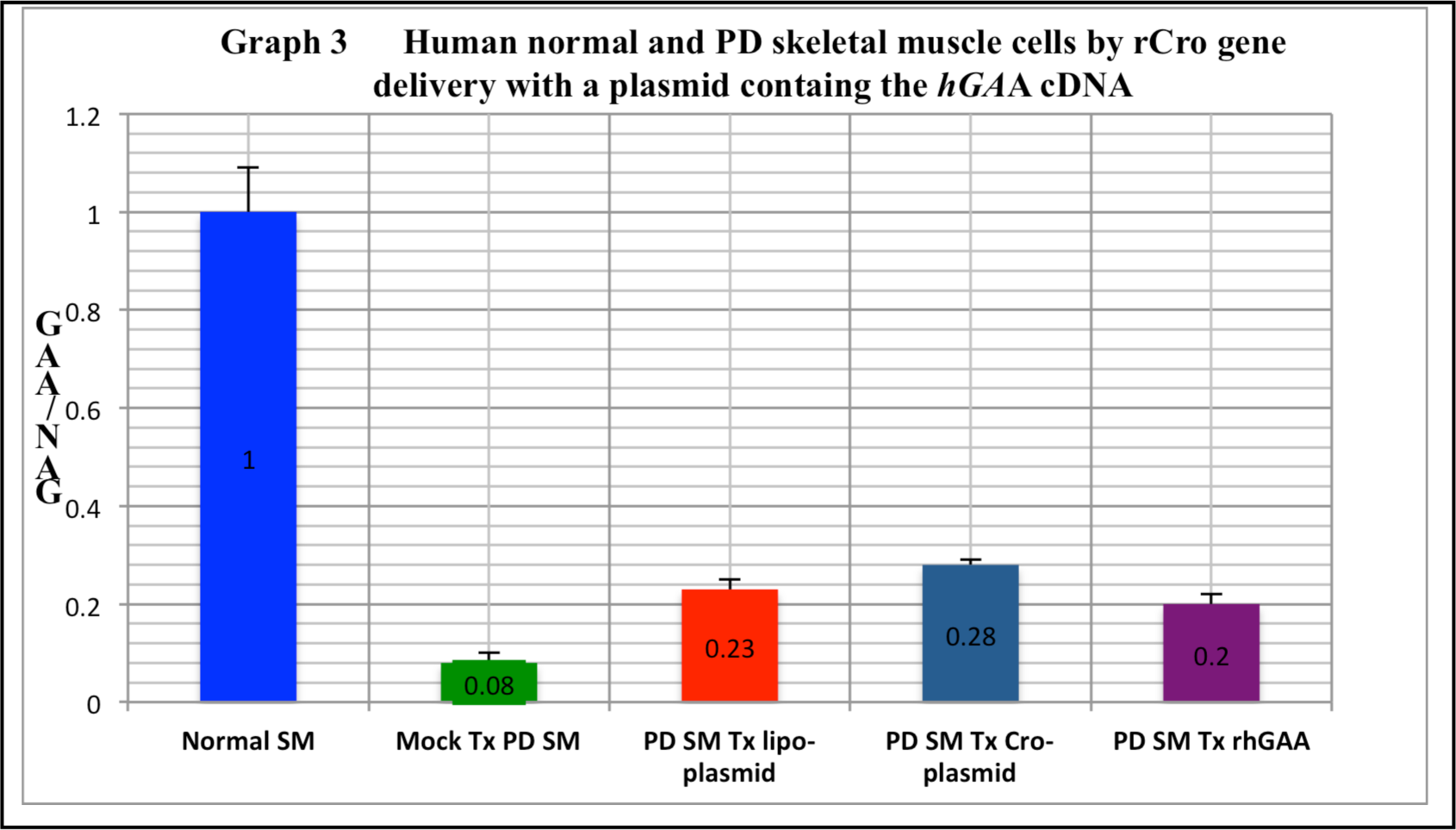
Human normal and PD myoblast cells by rCro-*hGAA* plasmid delivery. A human PD skeletal muscle cell line cell line was transfected with 4 µg of *hGAA* plasmid complexed to 10 µg of rCro or with Lipofectamine 3000-*hGAA* plasmid or exposed to rhGAA. After 48 hr, cells were assayed for GAA and NAG. Mock treated AMD and normal cell lines were controls. We found that lipo-*hGAA* plasmid increased GAA to 23% of normal while rCro-*hGAA* plasmid increased GAA to 28% of normal and rhGAA increased GAA to 20% of normal in PD myoblast cells (mean+/SD). All treatments were significant at p=≤0.05.

**Graph 4.**
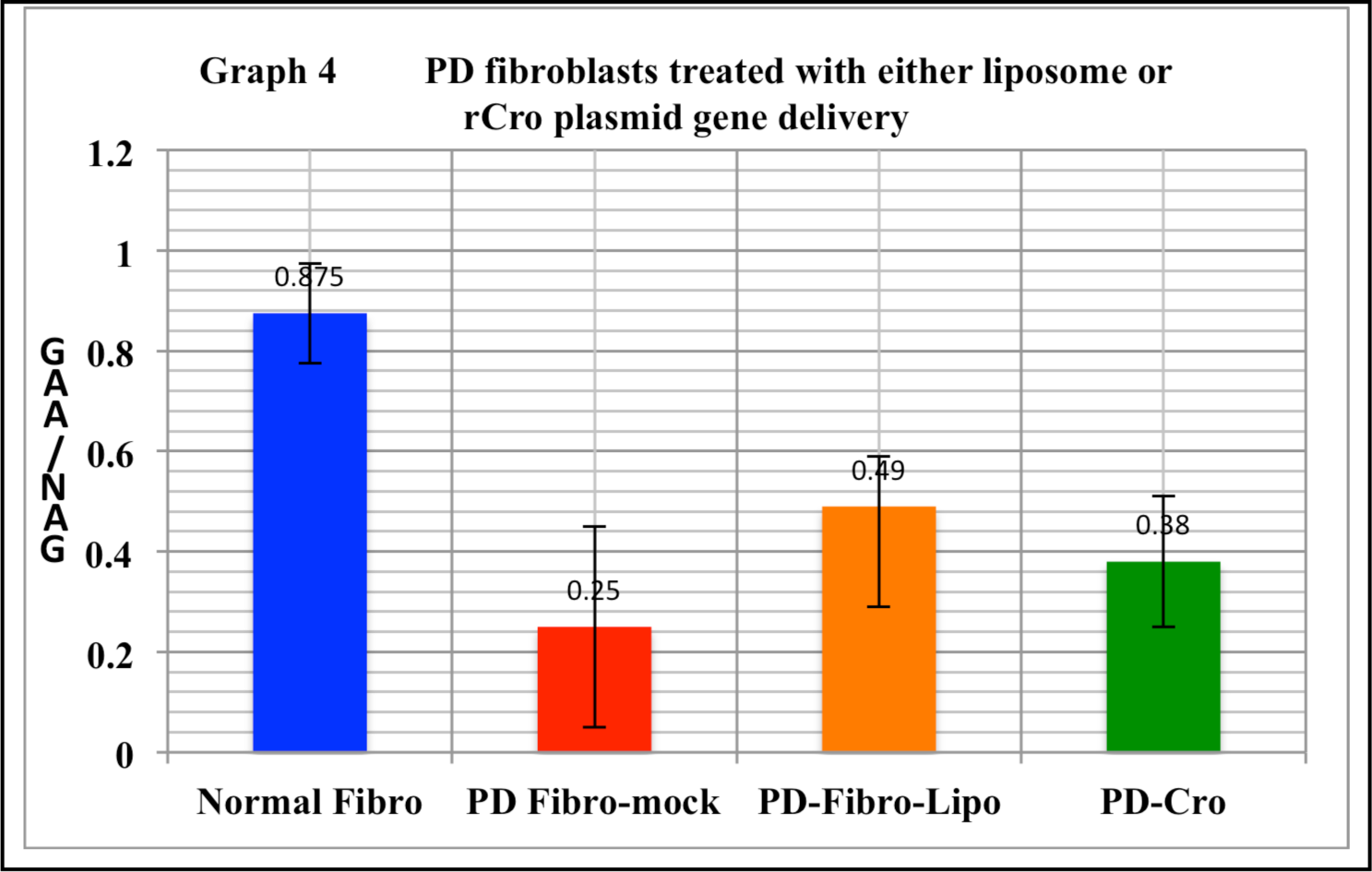
PD fibroblast cells treated with either liposome or rCro-*hGAA* plasmid delivery. Human three PD fibroblast cell lines were transfected with 4 µg of *hGAA* plasmid complexed to 10 µg of rCro or with Lipofectamine 3000-*hGAA* plasmid. After 48 hr, cells were assayed for GAA and NAG. Mock treated PD and normal cell lines were controls. In transfected PD fibroblast cell lines transfected with rCro-*hGAA* plasmid, increased GAA/NAG ratios was observed to various degrees as compared to normal cells. All treatments were significant at p=≤0.05.

**Graph 5.**
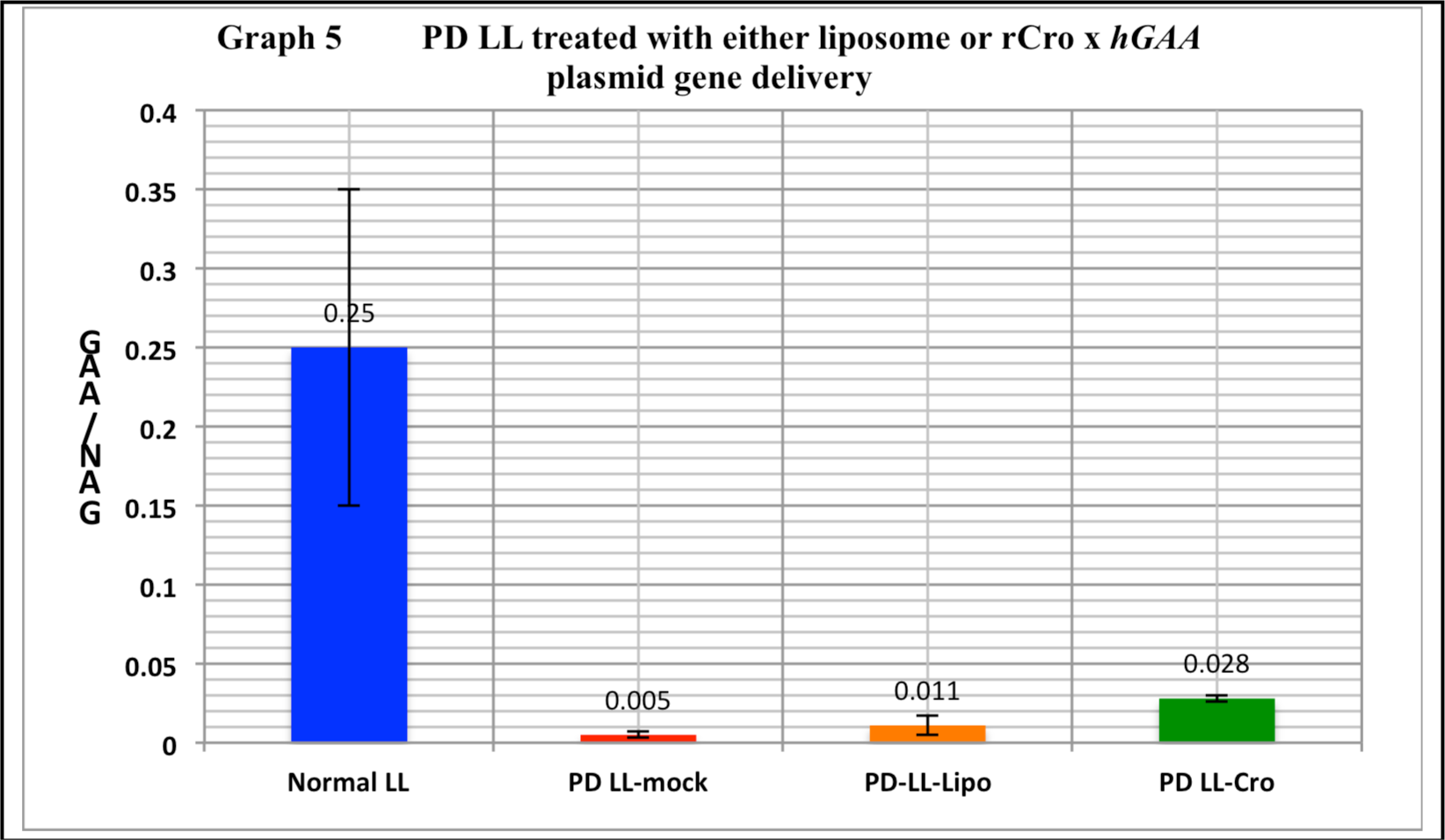
PD LL treated with either liposome or rCro-*hGAA* plasmid delivery. Human PD three lymphoid cell lines were transfected with 4 µg of *hGAA* plasmid complexed to 10 µg of rCro or with Lipofectamine 3000-*hGAA* plasmid. After 48 hr, cells were assayed for GAA and NAG. Mock treated AMD and normal cell lines were controls. In transfected PD lymphoid cell lines transfected with rCro-*hGAA* plasmid, increased GAA/NAG ratios was observed to various degrees as compared to normal cells. All treatments were significant at p=≤0.05.

